# Stitch-seq: Scalable CRISPR gene expression response profiling

**DOI:** 10.64898/2026.04.27.719216

**Authors:** Frances R. Keer, Aziz M. Al’Khafaji, Paul C. Blainey

## Abstract

Single-cell profiling of genetic perturbations has expanded our ability to map causal links between genes and phenotypes; however, the high cost and technical complexity of current methods restrict systematic interrogation of dynamic cellular programs. Here, we present Stitch-seq, a high-throughput pooled functional genomics sequencing method enabling simultaneous capture of CRISPR perturbations and targeted gene and protein expression across millions of cells. Stitch-seq utilizes single-cell droplet-based overlap-extension reverse-transcription PCR reactions to physically link gene expression features of interest to perturbation identifiers without cell barcoding or extensive sequencing. We validated Stitch-seq’s high fidelity using simplified models, benchmarked multi-omic Stitch-seq against single-cell RNA-sequencing in the MCF10A Epithelial-Mesenchymal Transition (EMT) model, and applied Stitch-seq to map transcriptional responses of MCF10A cells undergoing TGF-β-induced EMT to perturbations across five time points. By efficiently delivering large-scale multi-omic gene expression readouts, Stitch-seq provides a powerful and accessible modality for the routine dissection of complex biological pathways.

## Background

Defining the numerous complex and dynamic regulatory networks that govern cell state transitions remains a fundamental challenge in human biology and disease^1,2^. This problem has been the subject of intense interest in the context of cancer progression, where the cell regulatory state has been linked to patient prognosis^3^. Accurate mapping of regulatory networks requires large-scale molecular profiling capable of capturing shifts in the expression of key transcripts and proteins across many perturbations and time points– a capability that exceeds current methodologies.

Genetic perturbation screens, particularly those leveraging CRISPR technology, are powerful tools to establish causal links between genes and phenotypes. Historically, large-scale pooled perturbation screens have relied on enrichment-based phenotyping assays followed by bulk next-generation sequencing (NGS) to measure the relative abundance of perturbations in an enriched population^4–7^. While highly scalable, requiring only a single short-read NGS run and a simple statistical analysis, these approaches typically rely on univariate readouts (e.g., cell fitness^4,5^, differential binding of fluorescent antibodies or probes^7,8^, or specialized reporter systems^9^) that serve as simplified proxies for complex cellular processes. The constrained relevance and fidelity of enrichment screening data limit their utility for mapping and characterizing complex gene regulatory phenomena.

The integration of pooled perturbation libraries with single-cell RNA sequencing (scRNA-seq) readout, pioneered by Perturb-seq and CROP-seq^10–13^, offers high-dimensional– also known as “high-content”, “rich”, or “profiling”– molecular readouts of perturbation responses. In contrast to enrichment screens, high-dimensional readouts provide tremendous potential for discovery as the resulting datasets capture many distinct cellular and molecular phenotypes and permit post-hoc definition of target phenotypes and associated discoveries^10–19^. As such, these readouts yield rich information about the causal effects of gene perturbations on complex cellular programs, including cancer progression^20^.

Despite this technological advance, single-cell ‘omic profiling remains costly at the high cellular scale (10^5^-10^6^ cells)^19,21,22^ necessary to power large functional genomic screens. Consequently, most single-cell transcriptional profiling efforts have been severely limited in scale, typically using pooled CRISPR libraries with fewer than 100 gRNAs and only at one time point^20,23^, even in light of targeted scRNA-seq approaches and emerging approaches for sequencing cost reduction^24–27^. In spite of the high costs for single-cell-resolved profiling, several groups are embarking on large-scale Perturb-seq/CROP-seq data generation efforts at high expense^28^ in order to train “virtual cell” foundation models^29,30^. The push towards generating high-content perturbational data for such large efforts is a clear demonstration of the field’s conviction in the power of this data type for discovery and inference. However, the field lacks the capability to routinely enable large-scale profiling screens. To systematically dissect complex regulatory programs, a new screening modality is required– one that offers the richness of gene expression profiling with significantly improved cost efficiency compared with available single-cell ‘omics methods.

To address this need, we developed Stitch-seq, a perturbation profiling method that achieves the scalability necessary for the routine generation of large-scale high-content screening data. Stitch-seq profiles the response of a targeted set of mRNA- and protein-level gene expression features to perturbations in the large cell populations necessary for strong statistical power and perturbation effect quantification. Here, we demonstrate 11-plex multi-omic feature readout across a library of 120 perturbations with direct comparison to single-cell RNA sequencing with CITE-seq integration. We further demonstrate a 6-plex mRNA feature readout with inputs of approximately 800,000 cells per time point across five time points, totaling approximately 4 million cells processed in single-cell stitching reactions.

The Stitch-seq protocol leverages the unique thermostable reverse transcriptase, RTX^31^, combined with droplet-based overlap-extension PCR^32^ to physically link an expressed barcode or gRNA sequence to a user-defined multiplexed set of gene expression features (native mRNA and/or protein barcode sequences) within each single-cell droplet. This targeted “stitching” eliminates the need for specialized single-cell barcoding reagents and reduces sequencing depth requirements by orders of magnitude compared to Perturb-seq/CROP-seq. With Stitch-seq, gene expression counts and their associated perturbations are efficiently tallied for the targeted features via short-read amplicon sequencing, enabling straightforward computational demultiplexing of perturbation effects on the selected gene expression features. Stitch-seq occupies a unique and attractive position among available screening technologies, enabling molecular profiling screens that span more perturbations and conditions at lower costs.

Here, we report and evaluate the Stitch-seq protocol, demonstrating its utility by applying Stitch-seq to map the dynamic expression response of key Epithelial-Mesenchymal Transition (EMT) factors to CRISPR perturbations in the MCF10A system. EMT is a fundamental morphogenetic program characterized by a highly plastic continuum of cell states that drives cancer metastasis and therapeutic resistance^33^, with well-described markers delineating canonical epithelial-leaning or mesenchymal-leaning cell states. Investigating the regulatory mechanisms of transient states throughout this transition requires a profiling method capable of large cellular throughput and faithful multiplexed perturbation-readout association, such as Stitch-seq. In our large-scale EMT screen, Stitch-seq accurately recapitulates perturbation-specific cellular responses, validates canonical EMT regulators, and uncovers insights into the complex feedback loops governing cellular plasticity.

## Results

### Stitch-seq enables accurate perturbation profiling

Stitch-seq (Fig. 1a) physically links expressed perturbations of interest (e.g., a guide RNA) to a multiplexed panel of native mRNA and protein (via oligo-conjugated antibodies) targets, or features, in a droplet-based overlap extension reverse transcription single-cell PCR reaction. Sample preparation for Stitch-seq consists of staining cells with oligo-conjugated antibodies targeting surface proteins of interest^34^ and lightly fixing with paraformaldehyde (PFA). Next, we emulsify the cells in an overlap extension PCR mix using a custom two-inlet flow-focusing microfluidic chip with pressure controls (Supplementary Fig. 1a). We then thermocycle the droplets to selectively reverse transcribe the guide RNA (gRNA) and mRNA transcripts of interest, amplify the resulting cDNA and protein barcodes to add a common overlap sequence to each, and ultimately physically link the gRNA transcript to the feature amplicons via the complementary overlap sequence. The resulting “stitched” amplicons are removed from the emulsion and sequence library construction is completed via a nested qPCR to prevent overamplification. Following paired-end sequencing, we computationally demultiplex the mRNA transcript and protein-associated oligo counts on the physically linked perturbation sequence.

**Figure 1.**
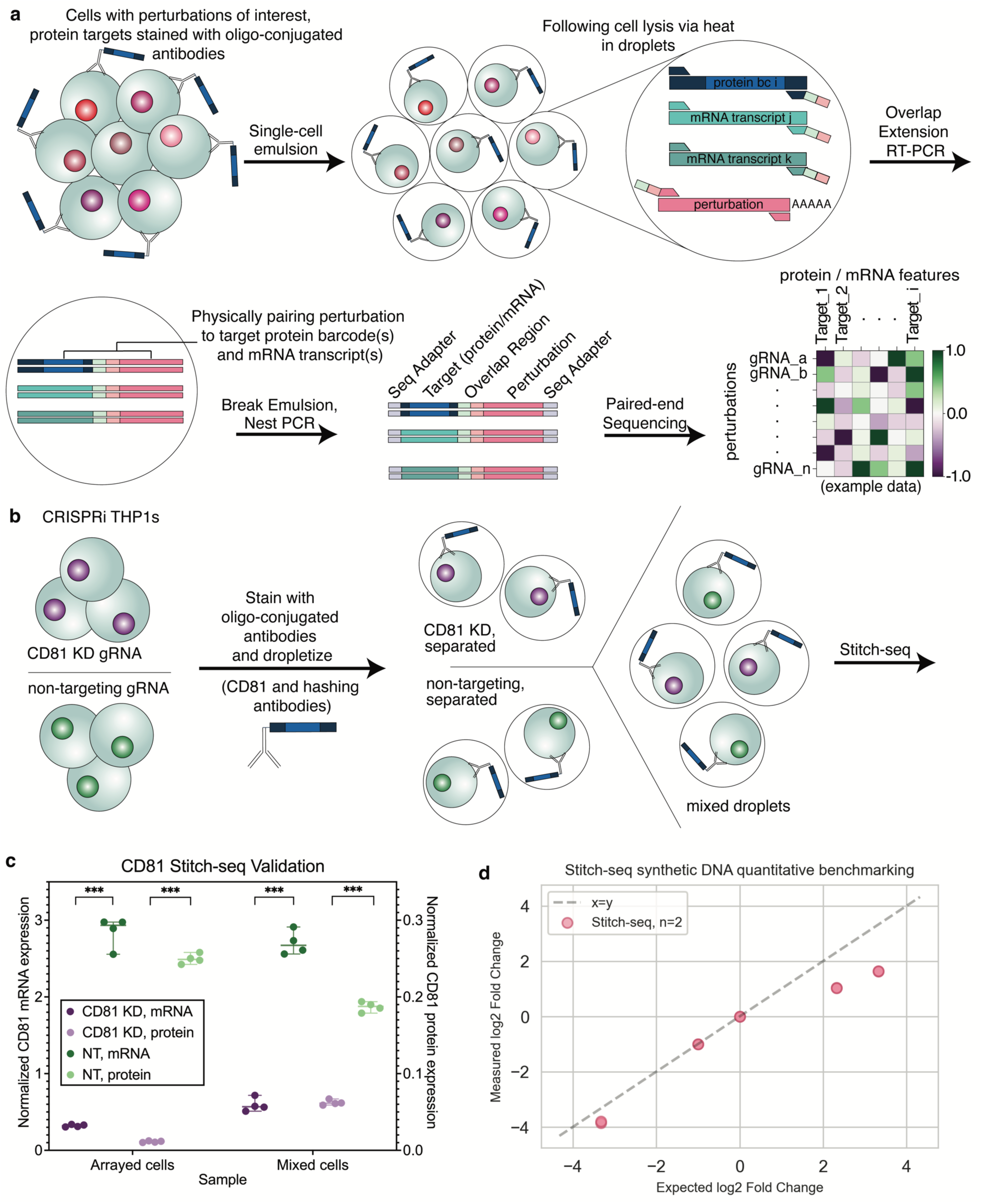
Stitch-seq enables accurate multi-omic perturbation profiling. **a**, Schematic overview of the Stitch-seq workflow. Cell populations transduced with a pooled CRISPR library are emulsified into single-cell droplets with overlap extension RT-PCR reagents. Following thermal lysis and reverse transcription, the overlap extension PCR physically links gRNA transcripts to target cDNAs or antibody-conjugated oligo barcodes from the single cells. These linked amplicons are subsequently analyzed via short-read sequencing to assign multi-omic phenotypes to specific genetic perturbations. **b,** Generation of the CD81 CRISPRi validation model. THP1 cells expressing dox-inducible dCas9-KRAB were independently transduced with either a non-targeting (NT) control gRNA or a validated gRNA targeting *CD81*. **c,** Stitch-seq accurately captures multi-omic knockdown in pooled cell populations. Profiling of arrayed and pooled NT and *CD81*-knockdown (KD) cell populations reveals a large and statistically significant reduction of both CD81 mRNA and protein expression, confirming high-fidelity signal capture and minimal inter-droplet crosstalk in a complex cellular context. Statistical significance was calculated using multiple unpaired t-tests (*** *p* < 0.001, *n* = 4). **d,** Quantitative accuracy and dynamic range of Stitch-seq readouts. A titration of synthetic DNA across a physiological range of concentrations (2 to 200 fM) demonstrates a high correlation (Pearson *r* = 0.971, *n* = 2) between input molecules and Stitch-seq read counts, validating the method’s ability to accurately capture varying expression levels.

To assess the quantitative accuracy and dynamic range of Stitch-seq, we performed a titration experiment using five synthetic DNA templates across a 100-fold concentration range (2, 10, 20, 100, 200 fM). Following sequencing, we normalized the relative read counts of each DNA template to the 20 fM DNA sequence as an internal control. The results were highly reproducible, and we observed a consistent correlation between input concentration and Stitch-seq read abundance, although the method exhibited a compression of dynamic range. Specifically, higher concentration templates were moderately underrepresented relative to the initial input (Fig. 1d), but the method still exhibited a Pearson correlation coefficient of *r* = 0.971.

Expecting that the high reproducibility of Stitch-seq would support strong statistical inference on perturbations, we then sought to validate Stitch-seq using an arrayed CD81 CRISPRi knock-down system in THP1 cells (Fig. 1b). We first confirmed that a previously validated gRNA targeting *CD81* significantly decreased both CD81 mRNA and protein expression upon doxycycline induction compared to a non-targeting gRNA^35^ (Supplementary Fig. 1d, e). We then performed multi-omic Stitch-seq on a 1:1 pooled population of two cell lines established by transduction with these guide RNAs (CRISPRi THP1 with a *CD81* gRNA and CRISPRi THP1 with a non-targeting gRNA) to evaluate the maintenance of perturbation assignment fidelity throughout the workflow, particularly during the emulsion PCR and remainder of the library preparation steps. Following sequencing, CD81 mRNA and protein read counts associated with each gRNA were normalized to an internal standard (*GAPDH* counts for mRNA reads, hashtag antibody barcode counts for protein reads) to account for gRNA representation differences. We observed large and statistically significant (*p* < 0.001) decreases in CD81 mRNA and protein expression in the *CD81* knockdown condition compared to the non-targeting gRNA, both when the lines were maintained separately and when pooled (Fig. 1c).

### Stitch-seq screens recapitulate single-cell transcriptomic readouts and known regulators of EMT

We subsequently profiled the effects of genetic perturbations on the Epithelial-Mesenchymal Transition (EMT) pathway using Stitch-seq in the MCF10A non-malignant human breast epithelial cell line, which was previously established as a TGF-β-inducible EMT model^20,36^.

We performed pooled CRISPR knockout screens targeting 120 known regulators of the EMT pathway, with guide RNAs selected from a previously published MCF10A EMT screen^20^. Upon doxycycline induction in the presence or absence of TGF-β, we expected to observe distinct transitions between epithelial and mesenchymal phenotypic states. To rigorously benchmark Stitch-seq against commonly used and high-performing single-cell multi-omic gene expression profiling protocols, we quantified EMT using both multi-omic Stitch-seq (targeting key EMT mRNA transcripts and surface proteins) and single-cell RNA-sequencing with CITE-seq (10x Genomics), focusing on a panel of six key EMT-associated mRNA transcripts and five EMT-associated surface proteins (Fig. 2a).

**Figure 2.**
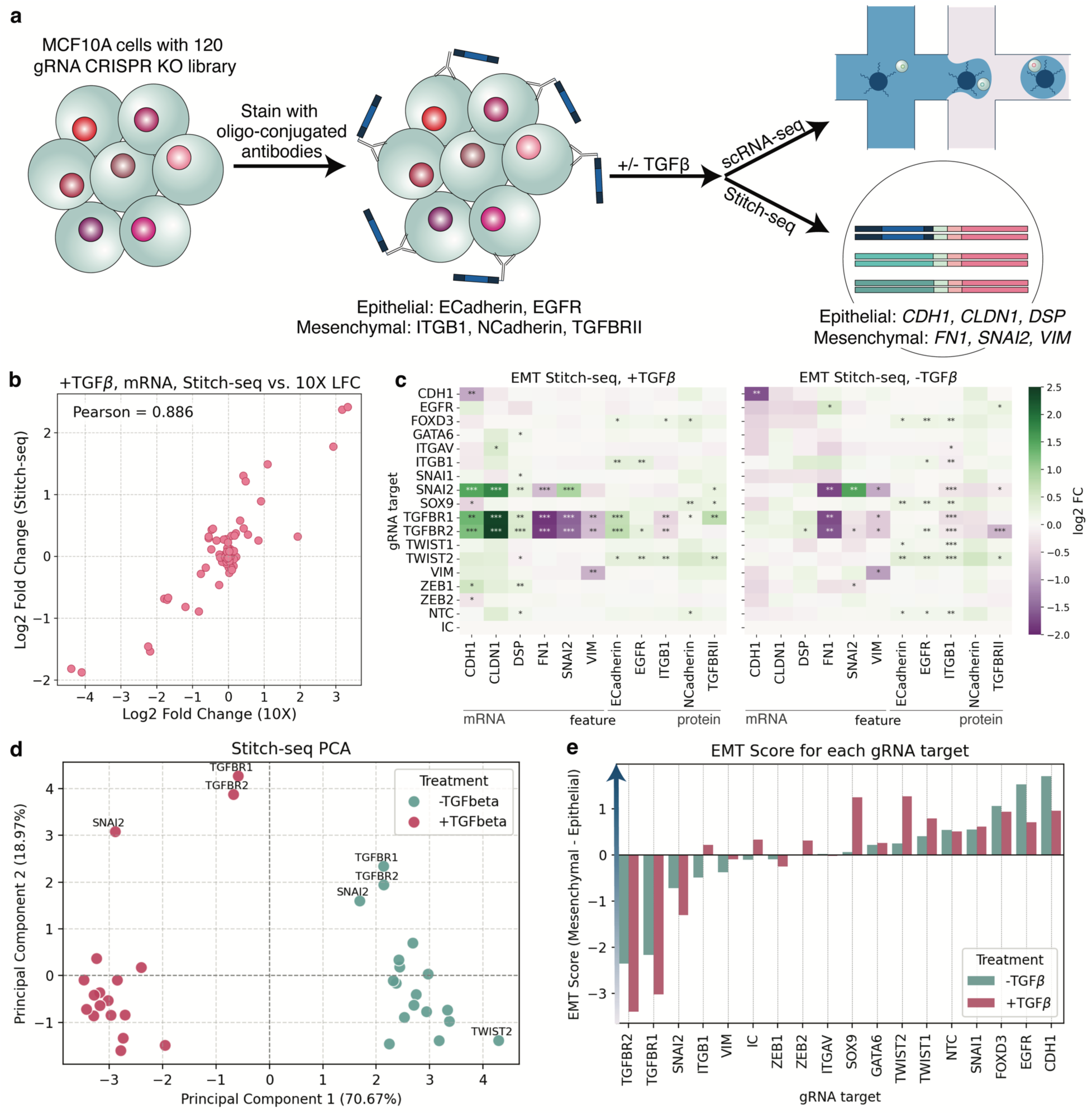
Stitch-seq screen identifies regulatory nodes and intermediate states in the EMT continuum. **a**, Parallel multi-omic profiling of EMT regulators. MCF10A cells were transduced with a 120 gRNA CRISPR knockout (KO) library targeting known and putative EMT regulators. Parallel profiling was performed using scRNA-seq integrated with CITE-seq (10x Genomics) and Stitch-seq to evaluate assay performance. **b,** Comparison of log2 fold changes (LFC) for targeted mRNA and protein features for every gRNA target and feature pair (here for +TGF-β mRNA condition: 6 mRNA targets x 18 perturbed genes = 108 pairs of LFC values, each of the 108 points is an average of *n* = 3 gRNA per perturbed gene values; see additional plots in Supplementary Fig. 2) reveals a Pearson correlation of up to 0.886 between Stitch-seq and single-cell RNA-seq LFC data, validating the correspondence of Stitch-seq and single-cell RNA-seq across diverse perturbations. **c,** Heatmaps of mRNA and protein expression shifts across gene perturbations +/- TGF-β. Data are normalized to *GAPDH* mRNA read counts, and fold change is calculated with respect to intergenic controls. Perturbations are averages of the 120 gRNAs across gene targets and heatmaps show 198 perturbation x readout values (average across *n* = 3). Significance was calculated via Mann-Whitney U-Test between the intergenic control population and each gRNA across the three replicates (* *p* < 0.05, ** *p* <0.01, *** *p* < 0.001). This reveals distinct functional modules of regulators that either promote or inhibit the transition to a mesenchymal state, including structural and feedback responses. **d,** Principal Component Analysis (PCA) of integrated mRNA and protein data. PCA was performed on the mRNA and protein expression data across the 18 targeted gene perturbations +/-TGF-β, showing distinct phenotypic clustering by treatment condition. **e,** EMT score analysis. Calculation of an “EMT score”, defined as the difference between the mean (average of *n* = 3) mesenchymal Z-scores (*FN1, SNAI2, VIM*, EGFR, ITGB1, N-Cadherin, TGFBRII) and epithelial Z-scores (*CDH1, CLDN1, DSP,* E-Cadherin), quantifies the global functional effect of each perturbation across the continuum.

For scRNA-seq, we achieved approximately 126,000 single cells per condition after computational demultiplexing and filtering, totaling 252,000 total cells profiled across two multi-omic screens. To directly compare performance between the two methods, we pseudobulked the scRNA-seq data by gRNA target and extracted the expression data corresponding to the 11 mRNA and protein Stitch-seq readouts.

Taking advantage of the high throughput of Stitch-seq, we then performed multi-omic Stitch-seq on three replicates of approximately 125,000 cells per replicate per condition, totaling 750,000 cells across six pooled multi-omic Stitch-seq gene expression screens. All read counts were normalized to *GAPDH* mRNA counts to account for differences in cell coverage for each gRNA. We subsequently collapsed the normalized reads for each feature by gRNA at the gene perturbation level. Perturbation effects were quantified as the log2 fold change (LFC) of each feature relative to the intergenic controls.

Comparison of the LFC values for the shared genes revealed a high correlation (Pearson correlation values between 0.45 and 0.9) between the two methods (Fig. 2b, Supplementary Fig. 2a). This demonstrates that Stitch-seq accurately quantifies the perturbation effects identified by conventional single-cell transcriptome-wide sequencing. Further, internal controls (knockout of genes encoding products in the readout set) verified these perturbations and Stitch-seq’s sensitivity, with targeted knockouts of *CDH1* and *VIM* yielding the expected reductions in their respective transcript levels (Fig. 2c). Additionally, the multi-omic capability of Stitch-seq captured decreases in ITGB1 and TGFBRII protein levels within their respective knockout populations, and significantly so in the absence of TGF-β (Fig. 2c). These internal controls verify the capacity of Stitch-seq to resolve perturbation-specific transcriptional shifts in individual gene products.

Beyond technical benchmarks, Stitch-seq successfully captured both canonical transcriptional regulatory networks and complex multi-omic feedback loops. Stitch-seq reported EMT patterns consistent with established literature (Fig. 2c), such as the knockout of the known *CLDN1* repressor, *SNAI2*, resulting in an increase in *CLDN1* expression (*p* < 0.001)^37^. Similarly, the loss of the signaling receptors, *TGFBR1*&*2*, led to an expected increase in *CLDN1* expression (*p* < 0.01, *p* < 0.001) by limiting cellular response to TGF-β, a known repressor of *CLDN1*^38,39^. Further, *SNAI2* knockout decreased *FN1* expression (*p* < 0.001), which is consistent with the previously observed positive relationship between the two genes^40^. Additionally, our targeted panel captured the activity of upstream E-box binding repressors, demonstrating that the depletion of *ZEB1* significantly upregulated *CDH1* (*p* < 0.05) and *DSP* (*p* < 0.01) transcript levels^41–43^.

With respect to proteins, under TGF-β treatment conditions, *TGFBR1*&*2* knockouts exhibited a significant upregulation of E-Cadherin protein compared to intergenic controls (Fig. 2c; *p* < 0.001)^44^. The retention of a core epithelial marker, E-Cadherin, in a normally pro-mesenchymal environment indicates that these perturbations limit initiation of EMT. Consistent with this blockade, while TGF-β treatment is a known potent stimulator of ITGB1 protein expression^45^, our results show that *TGFBR1/2* ablation effectively prevents this induction (*p* < 0.01), recapitulating the dependence of mesenchymal marker expression on intact TGF-β receptor signaling. Moreover, our panel captured elements of the canonical cadherin switch at the protein level, where depletion of the transcription factor SOX9 resulted in a significant upregulation of the mesenchymal adhesion marker N-Cadherin^46^ (*p* < 0.01), reinforcing its role in actively repressing mesenchymal adhesion programs. These findings recapitulate established regulatory relationships within the EMT pathway.

### Stitch-seq reveals complex multi-omic regulatory insights

Using Stitch-seq, we identified several previously uncharacterized multi-omic EMT regulatory relationships in the MCF10A model. Tracking protein and mRNA levels in tandem revealed compensatory surface protein remodeling in response to mechanical and structural disruption. For instance, the knockout of extracellular matrix receptor *ITGB1* triggered a significant upregulation of both EGFR and E-Cadherin surface proteins (Fig. 2c; *p* < 0.01 for both), suggesting that when cells lose integrin-mediated matrix attachment, they restabilize cell-cell junctions and upregulate epithelial growth factors to compensate. Similarly, depletion of the transcription factor *TWIST2* fundamentally altered the cell surface, driving significant increases in EGFR (*p* < 0.01), TGFBRII (*p* < 0.01), E-Cadherin (*p* < 0.05), and ITGB1 (*p* < 0.01) protein expression. We also observed evidence for receptor redundancy wherein *TGFBR1* knockout followed by TGF-β stimulation led to a significant increase in TGFBRII protein levels (*p* < 0.01).

At the transcriptional level, we identified *SNAI2* autoregulation within the EMT program. We observed a significant increase in *SNAI2* expression following *SNAI2* knockout in the absence of TGF-β treatment (Fig. 2c; *p* < 0.01), consistent with prior reporting^47^. Our findings further suggest a negative feedback mechanism whereby knocking out *SNAI2* activity by targeting CRISPR indels to the coding sequence (CDS) decreases repression of its own promoter, resulting in a net increase of *SNAI2* production. When cells were treated with TGF-β, we observed a slight dampening of this effect. Since TGF-β treatment is known to result in increased *SNAI2* expression, our findings suggest that the mechanism governing *SNAI2* self-regulation is altered by the presence of TGF-β signaling^48^.

To assess global phenotypic shifts, we performed Principal Component Analysis (PCA) on the integrated Stitch-seq mRNA and protein data. We observed separation based on TGF-β treatment along PC1, which comprised the majority of variance in the dataset (Fig. 2c, d). Additionally, knockout of either *TGFBR1* or *TGFBR2* caused a marked shift from the +TGF-β cluster towards the untreated control, confirming the loss of primary signal transduction. The *SNAI2* knockout mapped far away from both the epithelial and mesenchymal clusters, suggesting it plays a role in defining a partial EMT cell state^49–51^.

To distill these multi-omic readouts into a single metric reporting on epithelial-to-mesenchymal transition, we subtracted the mean Z-scores of epithelial markers (*CDH1, CLDN1, DSP*, E-Cadherin) from the mean Z-scores of mesenchymal markers (*FN1, SNAI2, VIM, EGFR*, ITGB1, N-Cadherin, TGFBRII) to calculate a composite “EMT score” (Fig. 2e) and summarize the molecular impact of each perturbation on EMT. This analysis identified *TGFBR1&2* and *SNAI2* as essential drivers away from the epithelial phenotypic state, with their loss resulting in a more epithelial-leaning phenotype, while *EGFR* and *CDH1* loss promoted transition. We further identified a class of genes, including *SOX9* and *TWIST2*, whose knockout sensitized cells to TGF-β-induced transition^46,52^. These knockouts showed negligible effects without TGF-β treatment but exhibited an exaggerated mesenchymal shift upon TGF-β exposure, suggesting that these factors may be critical for maintaining epithelial stability under environmental pressures.

### Stitch-seq scales to enable the high-density resolution of temporal dynamics of EMT

To demonstrate the scalability of Stitch-seq for high-resolution temporal profiling, we conducted 7-day time course experiments using the 120 gRNA CRISPR knockout library in MCF10As +/- TGF-β in triplicate, comprising an additional 30 gene expression profiling pooled screens. Following CRISPR induction, we split the population into TGF-β treated and untreated populations. Across days 1, 2, 3, 4, and 7 following the start of TGF-β treatment (with +/- TGF-β paired samples), we input a total of 4 million cells into Stitch-seq, targeting the key EMT mRNA markers of *CDH1*, *CLDN1*, *DSP*, *FN1*, *SNAI2*, and *VIM* (Fig. 3a). Further, this independently validated our previous day 7 results, showing high LFC correlation between the independent screens (Supplementary Fig. 3b).

**Figure 3.**
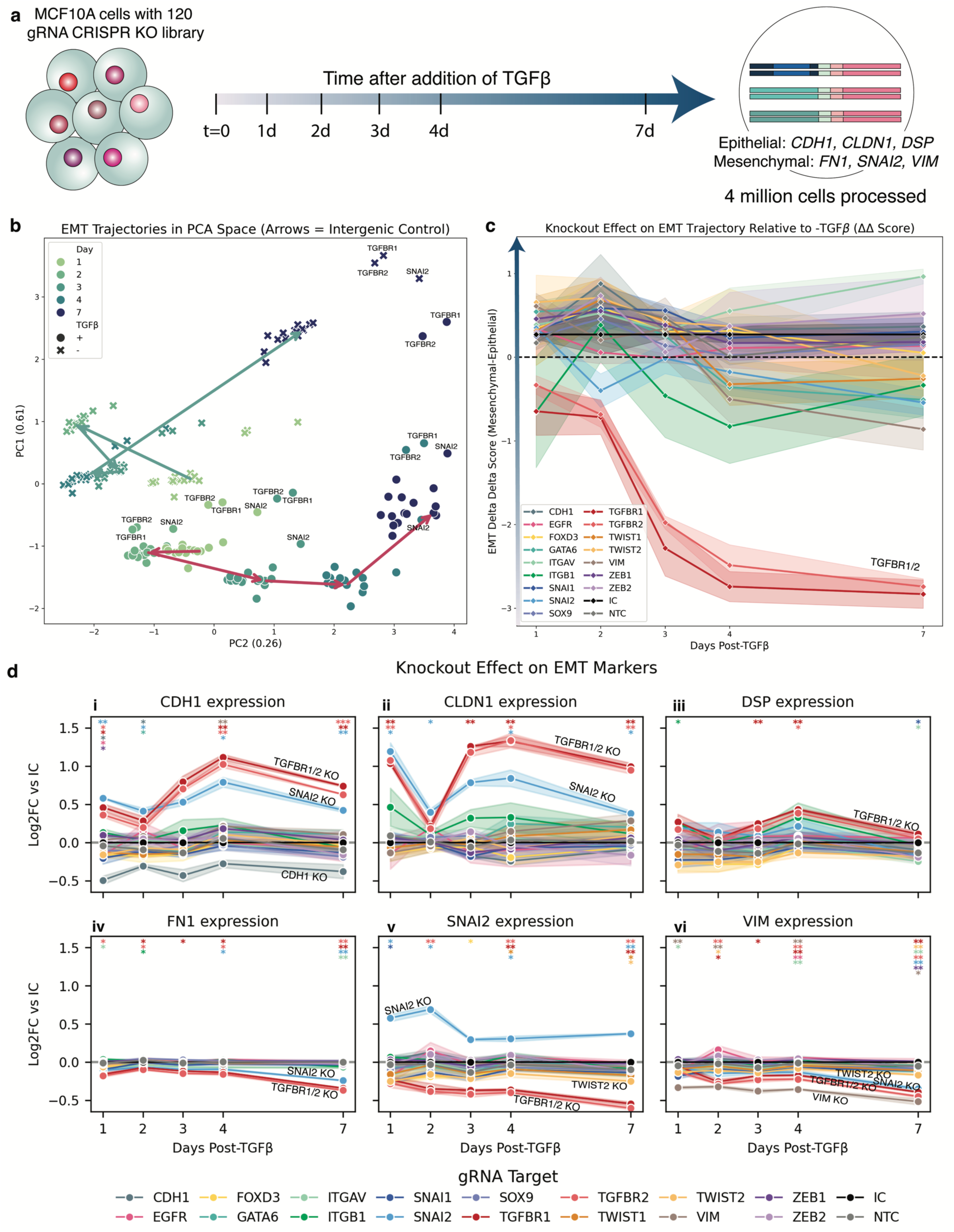
Large-scale Stitch-seq time point screen clarifies time dynamics of key mRNA transcripts throughout EMT. **a**, Longitudinal Stitch-seq profiling of EMT. MCF10A cells were transduced with a 120 gRNA CRISPR knockout (KO) library targeting known and putative EMT regulators. Cells were treated with doxycycline to induce Cas9 expression for 5 days. They were subsequently split into + TGF-β and - TGF-β conditions, and cells were collected at 1, 2, 3, 4, and 7 days post-treatment. Stitch-seq was then performed targeting key epithelial and mesenchymal mRNA markers. **b,** Principal Component Analysis (PCA) clustering of the Stitch-seq data across the time points and treatments. PCA shows distinct clustering by treatment condition and visualizes a continuous time trajectory for the +TGF-β population. Arrows represent the Intergenic Control trajectory. Data points represent a perturbation-gene expression pair. Loadings: *CDH1* 0.473, *CLDN1* 0.648, *DSP* −0.019, *FN1* −0.532, *SNAI2* −0.268, *VIM* −0.031. **c,** Composite “EMT ΔΔ Score”. Calculation of an “EMT ΔΔ Score” quantifies the global temporal effect of each perturbation. The score is defined as the difference between the mesenchymal (*FN1, SNAI2, VIM*) and epithelial (*CDH1, CLDN1, DSP*) Z-scores. These Z-scores were derived from double-normalized gene expression values: first, calculating the treatment effect as the log2 fold change of the +TGF-β values relative to the matched -TGF-β values (each as a mean of *n* =3), and subsequently, subtracting the corresponding treatment shift observed in the intergenic control population. **d,** Temporal dynamics of mRNA marker expression. LFC of mRNA marker expression compared to the intergenic control across the 7 days of TGF-β treatment shows that several gRNAs cause notable time-dependent changes in mRNA marker expression across the time points. Statistical significance is defined using a Benjamini-Hochberg False Discovery Rate threshold of 10% (* *q* < 0.1, ** *q* < 0.05, *** *q* < 0.01, *n* = 3 per time point for each perturbation/expression measurement pair).

To map the global trajectory of the transition, we performed PCA on the time course data (Fig. 3b). Mapping the biological progression of the intergenic control populations revealed a treatment-dependent divergence. While untreated cells maintained a relatively stable transcriptomic profile through day 4, the TGF-β treated cells exhibited a directed and continuous state transition along PC2 (loadings: *CDH1* 0.473, *CLDN1* 0.648, *DSP* −0.019, *FN1* −0.532, *SNAI2* −0.268, *VIM* −0.031) starting at day 2. Perturbations in *TGFBR1*, *TGFBR2*, and *SNAI2* caused distinct trajectory deviations. On day 4, these knockouts clustered with day 7 +TGF-β, but on day 7, the two *TGFBR* KOs clustered closer to the day 7 -TGF-β cells, indicating a dampened transition. Furthermore, the *TGFBR1&2* KOs from the TGF-β treated pools and the untreated pools clustered together.

To summarize complex multivariate transcriptional responses to TGF-β treatment into a single quantitative metric reflecting deviation from spontaneous EMT, we derived a composite “EMT ΔΔ score” (Fig. 3c). We first isolated the treatment effect by calculating the log2 fold change of the +TGF-β condition relative to the paired untreated samples. We then subtracted the corresponding log2FC of the intergenic control to isolate the targeting gRNA-specific changes. To quantify the transcriptional EMT shift, we finally calculated the difference of the mean Z-scores of epithelial markers (*CDH1, CLDN1, DSP*) from mesenchymal markers (*FN1, SNAI2, VIM*). Tracking this score over time unveiled the distinct transcriptional shift into an epithelial-leaning state for *TGFBR1&2* KOs by day 3.

To isolate the specific phenotypic consequence of each perturbation, we calculated the log2 fold change of each gene target relative to the matched intergenic control across the time points (Fig. 3d). Statistical significance was defined using a Benjamini-Hochberg False Discovery Rate threshold of 10% (FDR *q* < 0.1). This analysis revealed canonical EMT regulators and dynamic temporal phenomena.

Leveraging Stitch-seq’s temporal resolution, we captured both the early kinetics of epithelial dismantling and the temporal dependencies of intercellular junction remodeling. Consistent with the global PCA clustering results, depletion of *TGFBR1* or *TGFBR2* resulted in a significant failure to suppress *CLDN1* within 24 hours of TGF-β treatment (*q* = 0.023, *q* = 0.039) (Fig. 3d_iii_). Similarly, depletion of *SNAI2* induced a significant early retention of *CDH1* at day 1 (*q* = 0.017) (Fig. 3d_i_). These data confirmed that Smad-mediated activation of *SNAI2* (via *TGFBR1/2*) is required for EMT, as in its absence, key intercellular junction transcripts (*CLDN1*, *CDH1*) are upregulated within 24 hours^52^. However, continuous monitoring revealed that these dependencies are time-varying. Following the initial day 1 overexpression (*q* < 0.083), *CLDN1* levels under *TGFBR1/2* and *SNAI2* knockouts recovered on day 2, only to stabilize in an overexpressed state by day 7 (*q* < 0.073) (Fig. 3d_iii_).

From day 3 to day 7, the *TGFBR1*, *TGFBR2*, and *SNAI2* knockouts displayed substantially elevated *CDH1* and *CLDN1* expression. Knockout of these regulators induced a significant upregulation of the epithelial adhesion molecule *CDH1* (*TGFBR2 q* = 0.008; *TGFBR1 q* = 0.017; *SNAI2 q* = 0.046) (Fig. 3d_i_). The epithelial marker retention was mirrored by a concomitant failure to express characteristic mesenchymal genes, with *TGFBR1/2* and *SNAI2* knockouts all showing significant suppression of *FN1* and *VIM* at day 7 (*q* < 0.05) (Fig. 3d_ii_, Fig. 3d_vi_). We also observed that *TGFBR1* depletion resulted in the significant upregulation of *DSP* during the intermediate phases of transition at days 3 and 4 (*q* = 0.017, *q* = 0.038). Taken together, these data reflect the necessity of intact surface receptors for EMT pathway initiation^53^, while also demonstrating that ablation of downstream master regulators like SNAI2 renders cells incapable of approaching a mesenchymal-leaning transcriptional state^54^.

We also detected transient gene regulatory feedback and sequential transcription factor handoffs with our longitudinal transcriptomic profiling. In canonical EMT, *SNAI1* acts as an immediate-early activator of *SNAI2*^55^. Consistent with this, *SNAI1* perturbation resulted in a significant failure to upregulate *SNAI2* mRNA on day 1 (*q* = 0.099). Additionally, we observed a significant transient upregulation of *SNAI2* mRNA expression under the *SNAI2* knockout (Fig. 3d_v_), similar to our observations from our endpoint assay. This autoregulatory feedback loop initiated by day 1 post-treatment (*q* = 0.062), peaked at day 2 (*q* = 0.057), then decayed markedly (though ultimately remaining upregulated compared to the intergenic control) by day 3 and stabilized through day 7 (*q* = 0.024). This temporal signature further suggests a compensatory transcriptional feedback loop attempting to rescue *SNAI2* expression^47^.

Stitch-seq also uncoupled upstream regulatory networks from downstream structural effectors. To again validate the assay’s accuracy, we examined the *VIM* knockout (Fig. 3d_vi_). As expected, *VIM*-depleted cells showed a significant suppression of *VIM* mRNA across the time course, reaching maximal suppression by day 7 (*q* = 0.062). However, unlike the broad suppression of canonical EMT transcripts seen in the *TGFBR1/2* knockouts, *VIM* depletion triggered specific cytoskeletal feedback mechanisms^56^. On day 4, cells targeted for *VIM* knockout showed significantly upregulated *CDH1* (*q* = 0.023) (Fig. 3d_i_). The ablation of mesenchymal cytoskeletal proteins may compel the cell to transiently stabilize its epithelial adhesion networks, interfering with early and late-stage EMT^57^.

We further observed that depletion of *ITGAV* resulted in significant failure to maintain mesenchymal marker expression at the day 7 endpoint^58^, specifically impairing both *FN1* (*q* = 0.045) and *VIM* (*q* = 0.024) expression (Fig. 3d_ii_, Fig. 3d_vi_). Additionally, we identified transcription factor-specific cytoskeletal targets, such as *TWIST2*, which proved important for extended *VIM* expression at day 7 (*q* = 0.023) (Fig. 3d_vi_)^59^. These findings suggest that engagement with the extracellular matrix via integrin complexes likely provides signaling feedback that helps sustain the prolonged mesenchymal transcriptional program.

## Discussion

The landscape of CRISPR screening approaches for gene expression effects has been defined by a fundamental trade-off: high-throughput assays with unidimensional (enrichment) readouts or high-dimensional transcriptional profiling limited in throughput. Stitch-seq opens a new working regime by enabling high perturbation throughput with multiplexed multi-omic gene and protein expression readout. By physically linking gRNAs to transcripts of interest in single-cell reactions without single-cell barcoding, Stitch-seq circumvents financial barriers associated with existing single-cell whole-transcriptome sequencing workflows, substantially reducing the cost and time associated with multiplexed perturbation profiling in cases where phenotypes of interest can be captured in a targeted set of mRNA and protein features.

Critically, the development of Stitch-seq addresses the prohibitive cost and low throughput of standard single-cell technologies. By directly benchmarking our assay against Perturb-seq/CROP-seq with CITE-seq, we demonstrated that Stitch-seq faithfully recapitulates both transcriptional regulatory networks and dynamic surface protein landscapes. The robust correlation between Stitch-seq and single-cell RNA-seq validates our method as a reliable and highly scalable option for capturing complex multi-omic cell profiles.

By deploying Stitch-seq across a high-throughput 7-day TGF-β time course, we mapped the temporal perturbation response architectures in EMT. Typical single-endpoint screens are insensitive to early and intermediate dynamics and lose statistical power to call asynchronous expression effects when readout is only possible at one time point. Here we applied the throughput of Stitch-seq to cover multiple time points with replicate screens to show how transient cellular responses and feedback to perturbations can be reliably characterized. By reducing consumables cost and the sequencing effort to just 1% of standard whole-transcriptome workflows for similarly sized runs (Extended Data Table 1), Stitch-seq economically interrogated the complex temporal dynamics of the EMT response to TGF-β across 120 perturbations, 5 time points, and 4 million input cells.

The key distinction between Stitch-seq and Perturb-seq is the trade-off of single-cell resolution for scale, cost, and ease. While conventional Perturb-seq aims to capture the unique transcriptome of every individual cell, gene expression noise and technical factors often limit statistical power for hypothesis testing at the single-cell level. As a result, it is common practice to average single-cell screening data by condition, time point, and gRNA (pseudobulking) to achieve the statistical power necessary for perturbation effect calls and hit prioritization. Many analytical workflows go further to pseudobulk groups of cells clustered on a small subset of transcript targets that show good representation and variability across cells in a “feature selection” step. By contrast, Stitch-seq incorporates both pseudobulking and feature selection steps at the molecular level in the wet lab protocol, which drives the method’s relative simplicity and cost-efficiency. This efficiency enables the interrogation of 800,000 cells in one day—a scale that is currently cost- and time-prohibitive for whole-transcriptome single-cell approaches in most laboratories. Beyond its scalability in the perturbation dimension, the simplicity and low cost of Stitch-seq also permit the application of gene expression screens to complex experimental designs, spanning multiple cell lines, treatment conditions, and/or time points.

Building on this molecular efficiency, Stitch-seq’s cost reduction largely comes from its targeted capture of features of interest. In systems where *a priori* knowledge of key pathways is established, researchers often focus the interpretation of whole-transcriptome sequencing results on only a small fraction of the total transcripts. This mirrors other genomics applications, such as the shift in clinical oncology from untargeted Whole Genome Sequencing (WGS) to targeted DNA panels (e.g., for cancer diagnosis), which concentrate sequencing power on actionable mutations. While this same targeted philosophy has been applied to perturbation profiling, these methods still involve single-cell capture and initial cDNA preparation at the whole-transcriptome level, and as such, have been limited to less than 10-fold reductions in cost^24,25,60^ compared with whole-transcriptome scRNA-seq. Stitch-seq, however, achieves a 100-fold reduction (Extended Data Table 1) by capturing only features of interest in the generation of the sequencing library and by circumventing the need for expensive single-cell reagents. Additionally, while new hierarchical indexing strategies, such as 10x Flex Apex, enable dramatic cost reductions for cell preparation, such efficiencies are only achieved at massive cellular scale (Extended Data Table 1), and thus are unsuitable for routine and flexible use.

We envision Stitch-seq as a distinct and complementary approach to Perturb-seq/CROP-seq. Depending on the specificity of prior hypotheses about relevant expression phenotypes, users may choose to begin with more scale in perturbations (Stitch-seq) for systematic upstream discovery or more scale in expression readouts (Perturb-seq) for systematic downstream discovery. This enables a more flexible and project-specific approach to functional genomics, where researchers starting with Perturb-seq may want to expand discoveries on select expression phenotypes to additional perturbations, while others starting with Stitch-seq may want to expand select perturbations to additional readouts. Thus, we believe using a combination of Stitch-seq and single-cell perturbation profiling to perform discovery and investigation will be highly effective.

While Stitch-seq enables massive scalability in perturbation profiling thanks to its single-cell parallelized emulsion chemistry, the current protocol does not provide single-cell-resolved data outputs. Consequently, the assay cannot support analyses that require individual cell-quality filtering or tests for distributional shifts with invariant central statistics. However, this absence of single-cell resolution enables 1) vastly higher throughput than existing single-cell alternatives^10–13^ (permitting increased cell sampling), 2) the concentration of sequencing power exclusively on features of interest, and 3) the economic feasibility of independent replicate screens. Ultimately, this approach, producing data more similar to typical arrayed bulk RNA-seq applications, may result in superior overall statistical power under equivalent budget or time constraints. Beyond these analytical considerations, the protocols reported here were demonstrated up to an 11-plex readout. While higher-level multiplexing for Stitch-seq was not tested, this may require significant protocol modifications to overcome the limitations of multiplex PCR. Additionally, like many leading single-cell RNA-seq technologies^26,61,62^, Stitch-seq requires a large emulsion reaction to process cells. Fortunately, large-scale emulsion generation is a well-established practice in the genomics field and is achievable through a variety of approaches, including microfluidic devices (as shown here) or particle templating methods^63–66^.

The Stitch-seq framework described here provides a robust foundation for further development. Multiplexing capacity can be extended using larger panels of oligo-conjugated protein affinity reagents, and the protocol may be adapted to probe-based RNA-sequencing methods that circumvent the limitations of multiplex PCRs for heterogeneous transcripts. Additionally, incorporating clonal barcoding strategies may permit more complex experimental designs. Overall, Stitch-seq offers a distinct set of screening capabilities complementary to the current repertoire of CRISPR screening modalities and is well-suited for deeply characterizing the regulatory pathways that govern human health and complex disease states.

---

**Extended Data Figure 1.**
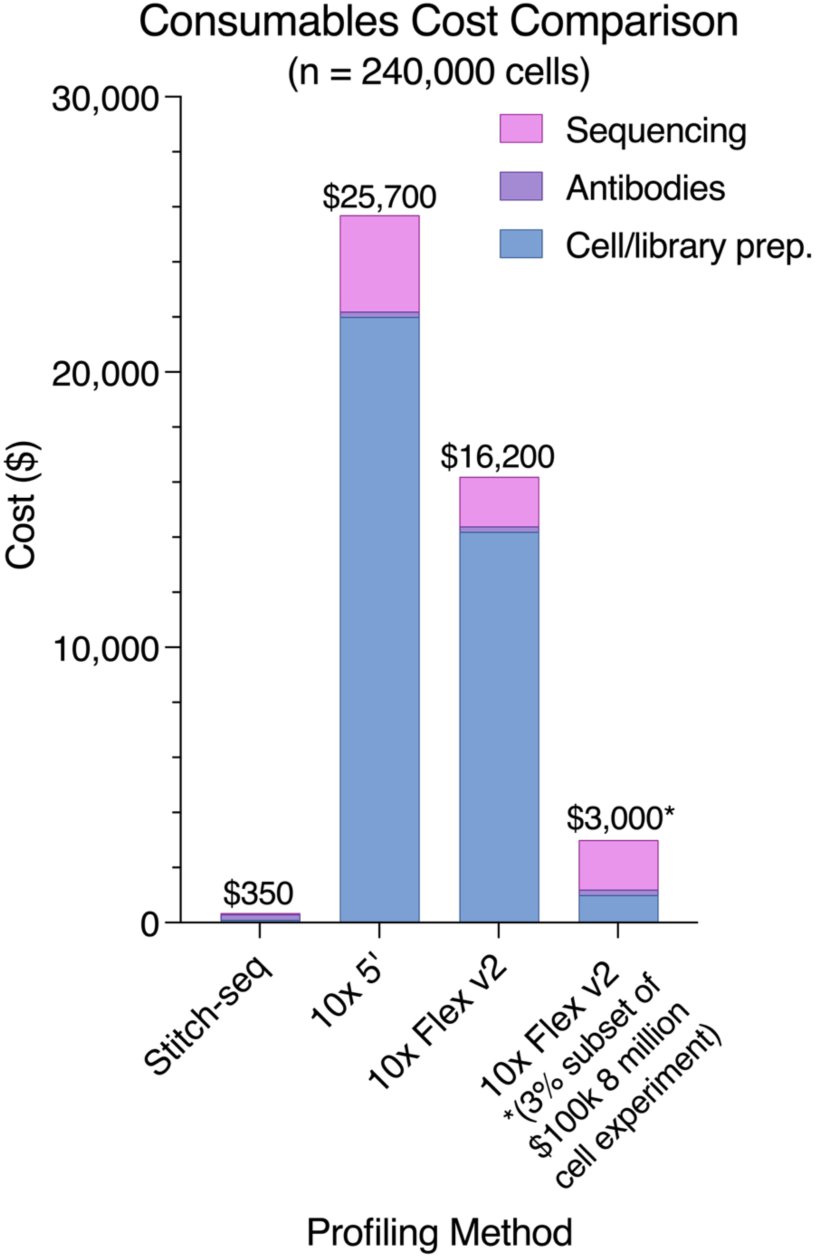
Visual comparison of consumables costs (as calculated in Extended Data Table 1) associated with 10x Genomics single-cell RNA-sequencing and Stitch-seq (n=240,000 cells)

**Extended Data Table 1.** Normalized comparison of 10x Genomics single-cell RNA-sequencing and Stitch-seq (n=240,000 cells)

| Item | Stitch-seq | Universal<br>scRNA-seq<br>(GEM-X Single<br>Cell 5' v3 kit) | Probe-based<br>scRNA-seq<br>(GEM-X Flex<br>v2 kit) | 240,000 cells<br>subset from a fully<br>multiplexed GEM-X Flex v2 chip |
| --- | --- | --- | --- | --- |
| Hardware cost<br>(one-time) | \$500<br>(customizable) | Requires<br>Chromium<br>Controller | Requires<br>Chromium<br>Controller | Requires<br>Chromium<br>Controller |
| Oligo-conjugated<br>antibodies (for<br>protein readout) | \$369 per Ab | \$369 per Ab | \$369 per Ab | \$369 per Ab |
| Cell/library prep<br>consumables cost<br>(240k cells) | \$100 + \$200 in<br>Ab<br>(5% of 10<br>antibodies) | \$22,000 + \$200<br>in Ab<br>(5% of 10<br>antibodies) | \$14,200 + \$200<br>in Ab<br>(5% of 10<br>antibodies) | \$1000<br>(3% of an 8<br>million cell run)<br>+ \$200 in Ab<br>(5% of 10<br>antibodies) |
| Total sequencing<br>effort | 75 million reads<br>(presented here) | 7.5 billion reads<br>(manufacturer<br>minimum) | 3.6 billion reads<br>(manufacturer<br>minimum) | 3.6 billion reads<br>(manufacturer<br>minimum) |
| Sequencing Read<br>Length | 300 cycles<br>(2x150) | 100 cycles | 100 cycles | 100 cycles |
| Approximate<br>Sequencing Cost<br>(NovaSeq X 25B) | \$50<br>(0.3% of 300<br>cycle kit) | \$3,500<br>(30% of a 100<br>cycle kit) | \$1,750<br>(15% of a 100<br>cycle kit) | \$1,750<br>(15% of a 100<br>cycle kit) |
| Total<br>consumables cost<br>for 240,000 cells | \$350 | \$25,700 | \$16,200 | \$3,000<br>(3% of a \$100,000<br>experiment) |

## Methods

### Quantitative benchmarking with synthetic DNA

Five synthetic DNA sequences of 200 bp were ordered, each with the same primer sequences (forward, outer, and inner) but with different internal sequences so they could be differentiated via sequencing. These were pooled at varying physiological concentrations (2, 10, 20, 100, 200 fM) and stitched, in bulk and in droplets, to a sixth synthetic DNA sequence representing the gRNA sequence-containing cDNA that would be present in a *bona fide* Stitch-seq reaction. The Stitch-seq protocol was completed and sequenced, and the resulting Stitch-seq read counts assigned to each synthetic DNA sequence were compared to the expected differences based on initial input concentration.

### CD81 CRISPRi model and benchmarking

Two gRNAs, one targeting *CD81* and one non-targeting, were cloned into array into the CROPseq-opti-vex vector, sequence verified, transfected into HEK293Ts with lipofectamine, and then transduced at MOI < 0.1 into THP1 cells engineered with a dox-inducible CRISPRi system. The cells successfully integrating a gRNA were selected using FACS based on GFP expression. CRISPRi expression was then induced with doxycycline for 3 days before the cells were assayed.

Bulk RNA-seq was performed on 1000 cells across 4 replicates per gRNA condition using the Smart-seq 2 protocol to confirm *CD81* mRNA expression knockdown. Flow cytometry was performed after staining with a PE anti-CD81 antibody (Supplementary Table 1) to confirm CD81 protein expression knockdown.

For Stitch-seq, cells were harvested, stained with CD81 and Hashtag 1 Total-seq B antibodies (Supplementary Table 1) according to the 10x Genomics Total-seq staining protocol, and fixed with 1% PFA (ThermoFisher #28906) at room temperature for 5 minutes. PFA was neutralized with a final concentration of 88 mM glycine solution. Cells were washed, resuspended at 0.8 ✕ 10^6^ cells/mL in 1X RTX buffer with 0.15% BSA (Supplementary Table 3), filtered with a 50 μm cell filter, and dropletized as two separate populations (CD81 and non-targeting) and as a 1:1 pooled population using a custom microfluidics system (Supplementary Fig. 1a) with Stitching PCR mix (Supplementary Table 2) and emulsion oil (Supplementary Table 4). Samples were then run through the full Stitch-seq protocol, sequenced, and analyzed to determine the number of CD81 protein and mRNA read counts that were physically linked to each gRNA relative to *GAPDH* mRNA counts and hashtag oligo barcode counts, respectively.

### Pooled EMT library design

gRNA sequences were collected from McFaline-Figueroa, et al.^20^, with a handful of additional gRNAs selected using the Broad Institute CRISPick portal^67^. These were ordered as an oPool and cloned into vector pRW494 using a golden gate reaction with a 1:6 ratio of backbone:insert. The plasmid library was then transformed into MC1061F-DUO electrocompetent cells (LGC 60514-1) such that we maintained 6,000 CFU per library member, before midi-prepping and performing a plasmid amplification qPCR and sequencing to confirm a 90/10 ratio under 3 with no library member dropouts. HEK293T cells were transfected with packaging plasmids and the library according to manufacturer protocols. The resulting virus was concentrated 100-fold using Lenti-X Concentrator (Takara #631231).

### MCF10A cell line engineering

MCF10A cells with dox-inducible Cas9 were transduced with varying concentrations of virus using a spinfection protocol, and puromycin selection was started 48 hours after transduction. On day 5 following transduction, samples with an MOI of less than 0.1 were selected and pooled such that a coverage of 1500 cells per library member was maintained.

### Pooled CRISPR screening for MCF10As

1.2 ✕ 10^6^ Cas9 MCF10As expressing the pooled CRISPR library were plated and treated with 1 ng/mL doxycycline (dox) for 5 days, with a passage and dox refresh at 2.5 days. After 5 days, the dox was discontinued and the cells were passaged into two flasks with 1.2 ✕ 10^6^ cells each, with or without 4 ng/mL freshly thawed TGF-β (ThermoFisher #PHG9214). TGF-β was refreshed at least every 48 hours for 7 days, with one passage at 3.5 days. Cells were visually assessed to confirm progression of EMT after 7 days, before being harvested.

### Single-cell transcriptomics for MCF10As

2.5 ✕ 10^6^ cells from each condition were treated for 15min with CellStripper (Corning #25-056-CI) and removed from the plate with a cell scraper, then were briefly treated with 1mL of TrypleE (ThermoFisher #12604013) for 2min followed by deactivation with media to decrease cell aggregation. The cells were washed and stained according to the 10x Genomics antibody staining protocol with a pool of Total-seq C antibodies (Hashtag: 0.5 μL, ITGB1: 0.5 μL, EGFR: 2 μL, E-Cadherin: 1.5 μL, TGFBRII: 3 μL, N-Cadherin: 3 μL) per sample (Supplementary Table 1), using optimal antibody amounts as previously determined by flow cytometry to maximize dynamic range. Following post-stain washing, cells were run through a superloading protocol for 10x 5’ RNA-seq with CRISPR and feature barcode (10x Genomics #PN-1000695). For superloading, 90,000 cells from each condition were loaded into 4 lanes of a 10x GEM-X chip, with the intent to demultiplex the doublets based on the presence of two or more gRNAs, such that each condition would yield data from at least 120,000 singlets. Library prep for the gene expression, protein, and CRISPR libraries were performed according to the 10x Genomics protocol, and the resulting libraries were sequenced on a NovaSeq X Plus 25B 100 cycle kit, with the cycle distribution 28/7/7/96.

### Single-cell transcriptomics data analysis

27 billion reads were run through CellRanger (Cell Ranger Multi v9.0.1) and the resulting files were further analyzed using custom scripts to filter out doublets as determined by the presence of more than one gRNA. Data was further filtered on cell quality. In order to directly compare to Stitch-seq, all genes targeted in the Stitch-seq panel were extracted and pseudobulked by gRNA target. Log2 fold change (LFC) values for each pseudobulked gRNA target were calculated with respect to the pseudobulked intergenic control population.

### Stitch-seq protocol (for MCF10As)

2.5 ✕ 10^6^ cells from each condition were treated for 15min with CellStripper and removed from the plate with a cell scraper, then were briefly treated with 1mL of TrypleE for 2min followed by deactivation with media to decrease aggregation. The cells were washed and stained according to the 10x Genomics antibody staining protocol with a pool of Total-seq B antibodies (Hashtag: 0.5 μL, ITGB1: 0.5 μL, EGFR: 2 μL, E-Cadherin: 1.5 μL, TGFBRII: 3 μL, N-Cadherin: 3 μL) per sample (Supplementary Table 1), using optimized volumes determined previously. Following post-stain washing, cells were fixed with 1% PFA at room temperature for 5 minutes, followed by deactivation with a final concentration of 88 mM glycine solution and subsequent dilution with PBS. Fixed cells were then washed twice with 1X RTX buffer (Supplementary Table 3) before ultimately being resuspended to a concentration of 0.8 ✕ 10^6^ cells/mL in RTX buffer with 1.5 mg/mL BSA and filtered with a 50 μm cell filter.

The stitching PCR mix was set up according to Supplementary Table 2, with primers for the Total-seq B antibodies, the gRNA, and the mRNA panel (*CDH1, CLDN1, DSP, FN1, SNAI2, VIM, GAPDH*). The stained and fixed cells, Stitching PCR mix, and Stitch-seq partitioning oil (Supplementary Table 4) were run through a custom droplet maker (Supplementary Fig. 1). The droplet maker was run for 30 minutes to collect 750 μL of droplets, using cell pressure of 0.5 psi, PCR buffer pressure of 1.8 psi, and oil pressure of 2 psi. Droplets were then gently washed with 500 μL of fresh partitioning oil three times before being aliquoted into PCR strip tubes with 20 μL of fresh partitioning oil and 20 μL of droplets. Droplets were transferred using the volume control of the micropipette instead of the pipettor button and with a wide-mouth pipette tip to minimize droplet shear stress.

PCR strip tubes were placed in a thermocycler with a preheated lid (105°C), set to a starting temperature of 20°C. The PCR reaction was performed according to the thermocycling conditions in Supplementary Table 5, with 2°C/sec ramping between every step.

After thermocycling, samples were removed and 100 μL of 20% 1H,1H,2H,2H-Perfluorooctanol in HFE7500 was added to each tube and incubated at room temperature for 5 min. 11 μL of aqueous phase from 12 PCR tubes was transferred to a fresh strip tube without collecting any oil phase. This was repeated two more times per condition for the other two replicates, such that each replicate should be pooled from approximately 130,000 cells. Following pooling, two SPRIselect (Beckman Coulter B23317) clean-ups were performed (0.8x and 0.7x) on each of the three replicates in the two conditions. All samples were ultimately resuspended in 20 μL of water.

### Stitch-seq library preparation

1 μL of each sample was added to two nested multiplex qPCR reactions, one for mRNA and one for protein, as detailed in Supplementary Table 6. Samples were thermocycled until amplification using cycling conditions described in Supplementary Table 8. Samples were 1x SPRI purified. 1 μL of each normalized sample was added to a P5/P7 qPCR reaction, as detailed in Supplementary Table 7, with each sample receiving a unique P7 primer. Samples were thermocycled until amplification using cycling conditions described in Supplementary Table 8. Samples were 1x SPRI purified, normalized via Qubit, and run on a Tape Station to determine average library member size before being pooled and sequenced with a 2×150 Element Aviti Cloudbreak FS mid-throughput sequencing kit (Element 860-00012).

### Stitch-seq data analysis

Paired end data was merged into a single-ended fastq using overlapping sequences with NGMerge for downstream processing. Data analysis was performed in a Python 3.7 environment. From fastq files, the gRNA primer binding sequences were removed using Cutadapt and the gRNA sequence was extracted and matched to a gRNA whitelist, with each read renamed to include its associated gRNA sequence. The known protein and mRNA overlap sequences were removed with Cutadapt and the remaining reads were aligned to a custom Bowtie2 reference genome for the protein or mRNA panel, respectively. Subsequent bam files were run through a custom Python script that counts alignments of each gRNA to each target gene. The count matrices were normalized to *GAPDH* counts, and the normalized gene counts assigned to each gRNA were compared. Significance was calculated via Mann-Whitney U-Test of the log2 fold change compared to the intergenic control across the three replicates.

### Stitch-seq primer design

Public mRNA sequencing data for MCF10As were downloaded from the Sequence Read Archive (SRA experiment accession: SRX16373062), aligned using Minimap2, and consensus sequences for each gene of interest were extracted. The longest protein coding sequence (CDS) was identified and input into Primer3Plus to pick 10+ fwd/rev primers with internal oligos such that the fwd/rev Tm is 63°C, internal Tm is 60°C, and fragment size is 180 bp with an internal oligo that is approximately 120 bp from the forward primer (Supplementary Fig. 1f). Those primers were then input into the Thermofisher multiple primer analyzer to check for primer dimers before being loaded into Geneious Prime to adjust Tms and lengths. The primers were then loaded into R and a modified TAP-seq pipeline^24^ was used to blast the primers and check for multiplex PCR compatibility. The primers were ultimately tested by performing a bulk Stitch-seq protocol on extracted mRNA and analyzing the products with custom scripts to check for mispriming and multiplex compatibility. All primers are provided in Supplementary File 1.

### Stitch-seq protocol for time point collection

Samples were collected for +/- TGF-β conditions on days 1, 2, 3, 4, and 7. For each time point, plated MCF10A cells (T-75) were washed twice with warm PBS and then treated for 15min at 37°C with 5mL of warm CellStripper. 1.5mL of warm TrypleE was added for 2min at 37°C. Cells were then gently removed from the plate with a cell scraper, followed by deactivation with 6 mL of high-FBS media to decrease cell aggregation. The cells were then resuspended in 5 mL of PBS with 1% BSA to stabilize the membranes. 2.5 ✕ 10^6^ cells were transferred to a new tube and lipid-modified oligos (LMOs, Sigma Aldritch LMO001) were then added at 1 reaction volume per 1 ✕ 10^6^ cells. The cells were pelleted and resuspended in fresh PBS then immediately stained according to the 10x Genomics antibody staining protocol (including FcX blocking) with a pool of Total-seq B antibodies (Hashtag: 0.5 μL, ITGB1: 0.5 μL, EGFR: 2 μL, E-Cadherin: 1.5 μL, TGFBRII: 3 μL, N-Cadherin: 3 μL, mouse isotype control: 0.5 μL, rat isotype control: 0.5 μL) per sample (Supplementary Table 1), matching previously optimized amounts. Following post-stain washing, cells were fixed with 1% PFA at room temperature for 5 minutes, followed by deactivation with 88mM glycine solution and dilution with PBS. Fixed cells were then washed twice with 1X RTX buffer with 1.5 mg/mL BSA before ultimately being resuspended to a concentration of 0.7 ✕ 10^6^ cells/mL in RTX buffer with BSA and filtered through a 50 μm cell filter. The cells were immediately dropletized with Stitch-seq PCR buffer, and thermocycled/processed according to the aforementioned Stitch-seq protocol. While LMO and antibody staining was performed, these data were not used in this manuscript.

To strictly ensure robust gRNA performance across the temporal profiling experiment, gRNA performance was evaluated independently within every biological replicate and time point. Expression values for all measured markers were globally Z-scored to ensure uniform dimensionality. A gRNA was flagged as an outlier in a given condition if its Euclidean distance from the target group’s median consensus signature exceeded a threshold of 3.0 standard deviations. To account for technical variation and random dropout, gRNAs were only excluded from global downstream analysis if they exhibited discordant phenotypes in greater than 50% of all measured conditions (days x replicates). As such, 6 out of 120 gRNAs were classified as chronically defective and removed from the final datasets.

To quantify global phenotypic shifts across the time course, a composite “EMT ΔΔ score” was derived for each perturbation. First, the treatment effect was isolated by calculating the log2 fold change of each marker in the +TGF-β condition relative to the paired -TGF-β value. The matching intergenic control value was then subtracted from each value to baseline normalize. These values were then standardized via Z-scoring across the dataset. The final composite EMT score was defined as the difference between the mean Z-scores of the mesenchymal markers (*FN1*, *SNAI2*, *VIM*) and the mean Z-scores of the epithelial markers (*CDH1*, *CLDN1*, *DSP*). A positive score indicates a mesenchymal-shifted phenotype while a negative score indicates an epithelial-shifted phenotype.

To evaluate the statistical significance of perturbation effects, the LFC of each gRNA target relative to the matched intergenic control baseline was calculated for every biological replicate. A two-sided one-sample t-test was then performed on the replicate LFC values against a null hypothesis mean of 0. Tests were conducted independently for each measured marker across all time points and gRNA targets, requiring a minimum of two biological replicates per condition. To correct for multiple hypothesis testing across the perturbation library, raw p-values were adjusted using the Benjamini-Hochberg False Discovery Rate (FDR) method. Statistical significance for all temporal biological discoveries was strictly defined by an FDR threshold of *q* < 0.1.

## Acknowledgements

We would like to thank the following individuals for feedback on the manuscript: Anna Le, Robert Majovski, Adrian Paskert, and Andrew Aguirre. We would also like to thank the following individuals for helpful conversations throughout the project: Anna Le, Russell Walton, Mohamad Najia, Lexi Fernandez, Justine Shih, Juniper Elacqua, Guoping Wang, as well as all members of the Blainey Lab and Methods Development Lab. Russell Walton provided the MCF10A Cas9 cell line and the vector backbone for the EMT library. Russell Walton and Anna Le provided molecular cloning protocol support. Victoria Popic wrote the first version of two scripts used to analyze Stitch-seq data. Fadi Najm provided the CD81 knockdown gRNA sequence. We would like to acknowledge Rebecca Dertinger and Martina Enev for administrative support. F.R.K. is supported by the National Science Foundation Graduate Research Fellowship Program under Grant No. 2141064. Any opinion, findings, and conclusions or recommendations expressed in this material are those of the author(s) and do not necessarily reflect the views of the National Science Foundation. Research reported in this publication was supported by the National Cancer Institute of the National Institutes of Health under Award Number R21CA269103. The content is solely the responsibility of the authors and does not necessarily represent the official views of the National Institutes of Health. Research reported in this publication was supported by the Broad Institute of MIT & Harvard under the Scientific Projects to Accelerate Research and Collaboration (SPARC) program. We acknowledge the in-kind contribution of Chromium reagents from 10x Genomics used in this work.

## Author contributions

A.M.A. and F.R.K. conceived of the project. F.R.K designed and performed experiments, analyzed data, and wrote the manuscript. A.M.A. and P.C.B. supervised the work and edited the manuscript.

## Data availability

Data is available at Gene Expression Omnibus (GEO) under accession GSE325318.

## Code availability

Code is available at https://github.com/fkeer/Stitch-seq.

