## Supplementary Information for "Stitch-seq: Scalable CRISPR gene expression response profiling"

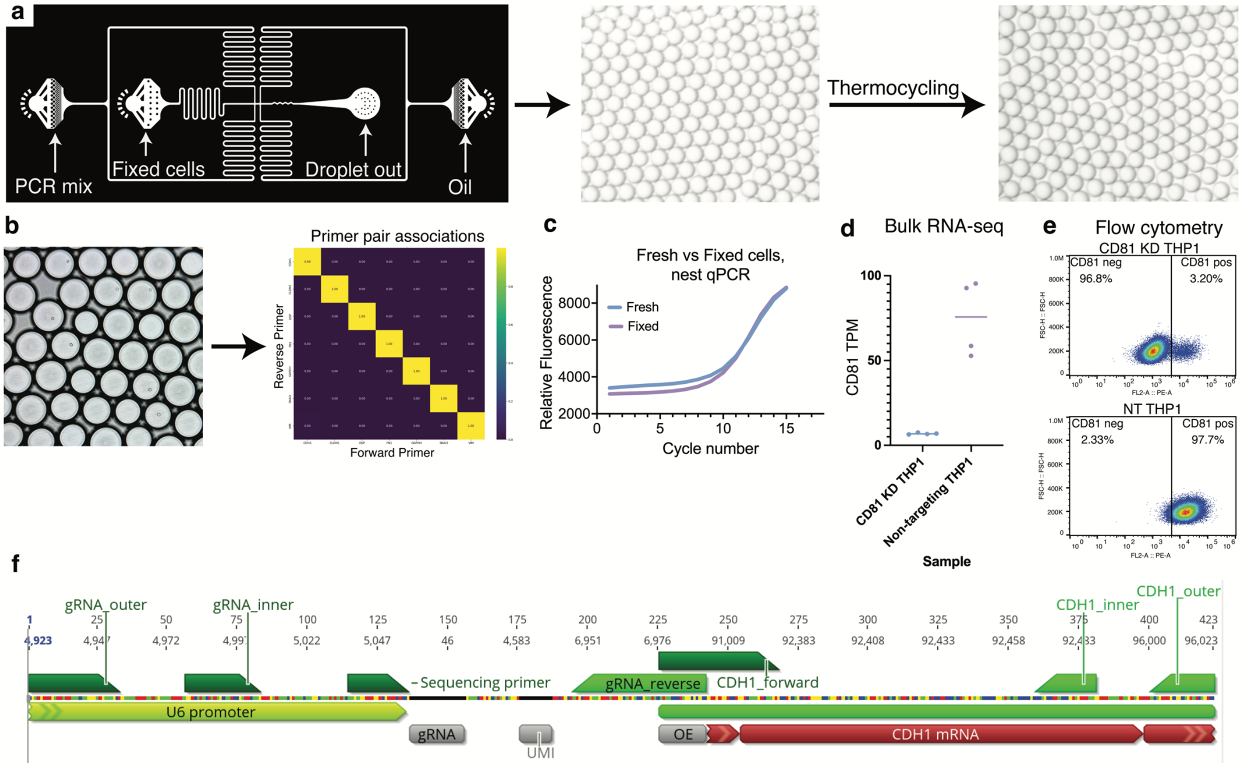


**Supplementary Figure 1 | Technical optimization, primer design, and orthogonal validation of the Stitch-seq platform.**

**a,** Emulsion generation and validation. Emulsions were generated using the depicted custom droplet-maker schematic. Optimization of the PCR buffer and fluorinated partitioning oil composition resulted in highly monodisperse droplets that successfully maintained compartmentalization throughout the physical stress of thermocycling.

**b,** Assessment of multiplexing fidelity. Single MCF10A cells were emulsified and subjected to the Stitch-seq protocol with the EMT mRNA primer panel. Sequencing readouts demonstrated negligible primer crosstalk during the 6-plex droplet RT-PCR.

**c,** Impact of cellular fixation on assay performance. Stitch-seq was evaluated in parallel on both fresh and 1% PFA fixed non-targeting KD CRISPRi THP1 cells, revealing negligible differences in CD81 transcript capture efficiency due to fixation.

**d,** Orthogonal transcriptomic validation of the CD81 CRISPRi knockdown model. Confirmation of significant *CD81* depletion at the transcriptional level in the transduced THP1 cells was achieved via bulk RNA-sequencing. This established a rigorous mRNA ground-truth of *CD81* depletion for subsequent Stitch-seq benchmarking.

**e,** Orthogonal proteomic validation of the CD81 CRISPRi knockdown model. Confirmation of significant CD81 depletion at the proteomic level in the transduced THP1 cells was achieved via flow cytometry. This established a rigorous protein ground-truth of CD81 depletion for subsequent Stitch-seq benchmarking.

**f,** Schematic representation of Stitch-seq primer design and overlap-extension mechanism. Each targeted feature utilizes an outer and inner nested primer pair. The forward primer for each target features a 5' overhang complementary to the reverse primer of the gRNA amplicon, enabling the physical stitching of the perturbation barcode to the phenotypic readout during the emulsion PCR.

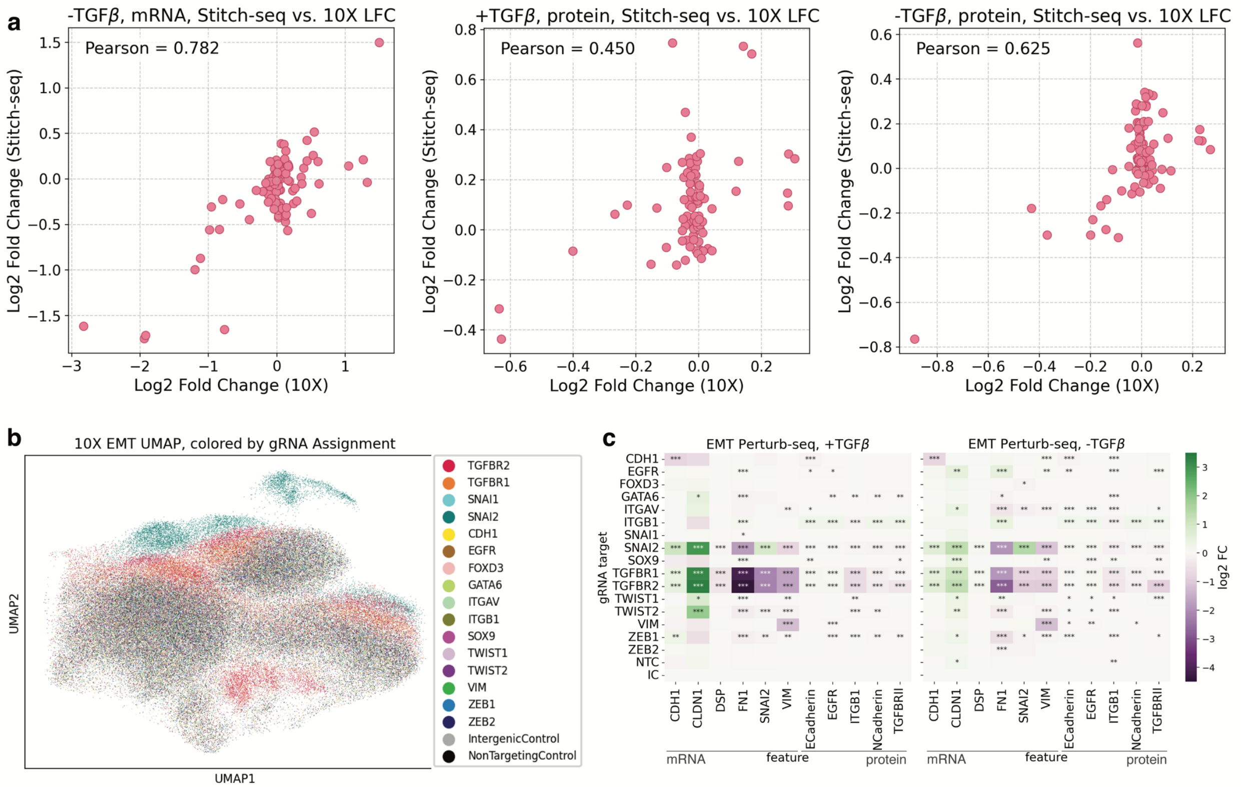


**Supplementary Figure 2 | Extended benchmarking of Stitch-seq against multi-omic single-cell sequencing.**

**a,** Concordance between Stitch-seq and transcriptome-wide readouts. Comparison of log2 fold changes (LFC) for targeted mRNA and protein features reveals a Pearson correlation between 0.450 and 0.886 between Stitch-seq and the scRNA-seq data.

**b,** UMAP analysis of single-cell RNA-seq performed on Day 7 of TGF-β treatment. Dimensionality reduction of the transcriptome-wide data visualizes the global phenotypic shifts resulting from perturbations.

**c,** Targeted analysis of scRNA-seq data. To enable direct benchmarking, the transcriptome-wide scRNA-seq data were comparably analyzed by extracting only the genes and proteins included in the Stitch-seq panel to identify significant mRNA and protein expression changes across the perturbations. Significance is determined by Mann-Whitney U-test on the differences in single-cell population gene expression compared to the intergenic control population.

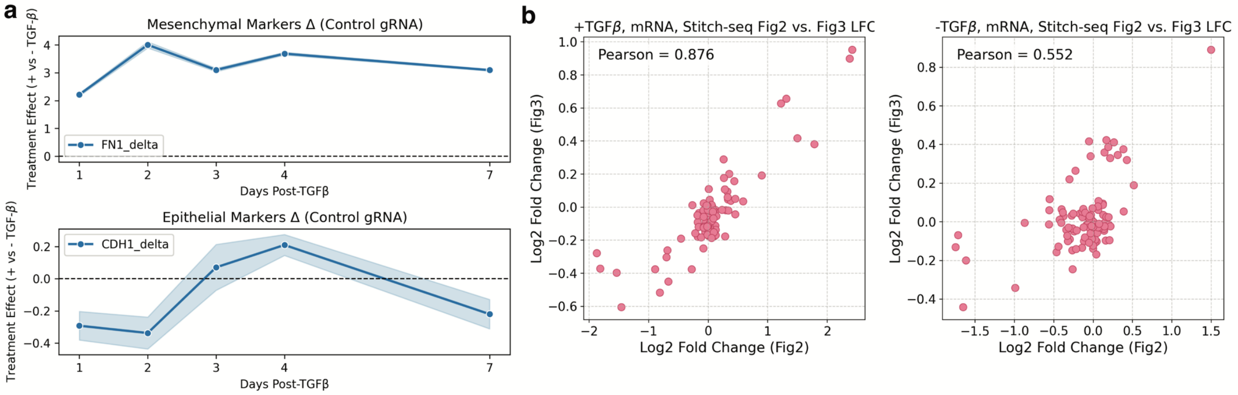


**Supplementary Figure 3 | Temporal expression dynamics and independent replicate concordance of the Stitch-seq time-course experiment.**

**a,** Baseline temporal kinetics of EMT markets. Expression differences of +TGF-β and -TGF-β conditions for *FN1* (mesenchymal) and *CDH1* (epithelial) within the intergenic control gRNA population demonstrate the expected induction of mesenchymal markers and suppression of epithelial markers, revealing distinct temporal kinetics across the 7-day transition.

**b,** Assay reproducibility across independent experiments. Comparison of the day 7 Stitch-seq data from the longitudinal time-course experiment against the day 7 mRNA data from the multi-omic benchmarking experiment demonstrates high concordance between the two independent Stitch-seq runs, validating the reproducibility of the platform.

**Supplementary Table 1: Antibodies with catalogue numbers**

| Antibody | Biolegend Cat. No |
|---|---|
| PE anti-human CD81 (TAPA-1) Antibody | 349505 |
| TotalSeq™-B0373 anti-human CD81 (TAPA-1) Antibody | 349525 |
| TotalSeq™-B0251 anti-human Hashtag 1 Antibody | 394631 |
| TotalSeq™-B0369 anti-human CD29 Antibody | 303033 |
| TotalSeq™-B0132 anti-human EGFR Antibody | 352929 |
| TotalSeq™-B0135 anti-human CD324 (E-Cadherin) Antibody | 324129 |
| TotalSeq™-B0433 anti-human CD325 (N-Cadherin) Antibody | 350825 |
| TotalSeq™-B1171 anti-human TGF-β Receptor II Antibody | 399719 |
| TotalSeq™-C0251 anti-human Hashtag 1 Antibody | 394661 |
| TotalSeq™-C0369 anti-human CD29 Antibody | 303029 |
| TotalSeq™-C0132 anti-human EGFR Antibody | 352931 |
| TotalSeq™-C0135 anti-human CD324 (E-Cadherin) Antibody | 324127 |
| TotalSeq™-C0433 anti-human CD325 (N-Cadherin) Antibody | 350819 |
| TotalSeq™-C1171 anti-human TGF-β Receptor II Antibody | 399717 |

**Supplementary Table 2: Stitch PCR mix for dropletization**

| Reagent | Final Concentration |
|---|---|
| gRNA outer primer | 0.5 μM |
| gRNA rev primer | 0.5 μM |
| mRNA panel fwd primer pool | 0.25 μM |
| mRNA panel outer primer pool | 0.25 μM |
| Protein fwd primer | 0.25 μM |
| Protein outer primer | 0.25 μM |
| RTX buffer | 2X |
| dNTPs | 0.4 mM |
| Betaine | 1.98 M |
| DTT | 4.88 mM |
| BSA | 1 mg/mL |
| RTX enzyme | 6.4 ng/μL |
| H2O | To 25 μL |

**Supplementary Table 3: 10X RTX buffer composition**

| Reagent | Final Concentration |
|---|---|
| Tris-HCl (pH 8.4) | 600 nM |
| $(NH_4)_2SO_4$ | 250 nM |
| KCl | 100 nM |
| $MgSO_4$ | 10 nM |
| H2O | To volume |

**Supplementary Table 4: Stitch-seq partitioning oil composition**

| Reagent | Volume |
|---|---|
| QX200™ Droplet Generation Oil for EvaGreen (BioRad #1864005) | 7 mL |
| 10% RAN-008 Fluorosurfactant (RAN 008-FluoroSurfactant-50G) in HFE-7500 (3M #ID7100025016) | 1.4 mL |

**Supplementary Table 5: Stitching PCR cycling conditions**

| Step | Temperature | Time | Cycles |
|---|---|---|---|
| Hold | 20°C | Until skip | 1 |
| Lyse | 80°C | 2 min | 1 |
| Reverse Transcription | 60°C | 2 min | 3 |
| | 68°C | 8 min | |
| Initial Denaturation | 94°C | 2 min | 1 |
| Denaturation | 94°C | 30 sec | 32 |
| Annealing | 60°C | 30 sec | |
| Extension | 68°C | 2 min | |
| Final Extension | 68°C | 7 min | 1 |
| Hold | 10°C | inf | 1 |

**Supplementary Table 6: Nest qPCR mix**

| Reagent | Final Concentration |
|---|---|
| gRNA inner primer | 0.5 µM |
| mRNA panel inner primer pool<br>OR<br>Protein inner primer | 0.5 µM |
| Platinum Taq II buffer | 1X |
| dNTPs | 0.2 mM |
| SYBR green | 1X |
| Platinum Taq II | 0.04 U/µL |
| Template | 1 µL |
| H2O | To 25 µL |

**Supplementary Table 7: Indexing qPCR mix**

| Reagent | Final Concentration |
|---|---|
| gRNA_P5 primer | 0.5 μM |
| R2i7P7 primer (different per sample) | 0.5 μM |
| Platinum Taq II buffer | 1X |
| dNTPs | 0.2 mM |
| SYBR green | 1X |
| Platinum Taq II | 0.04 U/μL |
| Template | 1 μL |
| H2O | To 25 μL |

**Supplementary Table 8: Nest/indexing qPCR cycling conditions**

| Step | Temperature | Time | Cycles |
|---|---|---|---|
| Initial Denaturation | 94°C | 2 min | 1 |
| Denaturation | 94°C | 30 sec | Until amplification |
| Annealing | 60°C | 30 sec | |
| Extension | 72°C | 30 sec | |
| Imaging | 72°C | image | |
| Final Extension | 72°C | 2 min | 1 |

**Supplementary File 1:** PCR primer sequences

**Supplementary File 2:** EMT library gRNA sequences

**Supplementary File 3:** Significance values from temporal analysis of Stitch-seq

**Supplementary File 4:** PCA values from temporal analysis of Stitch-seq
