## Supplementary figures and images for "Stitch-seq: Scalable CRISPR gene expression response profiling"

### Supplementary Figure 1

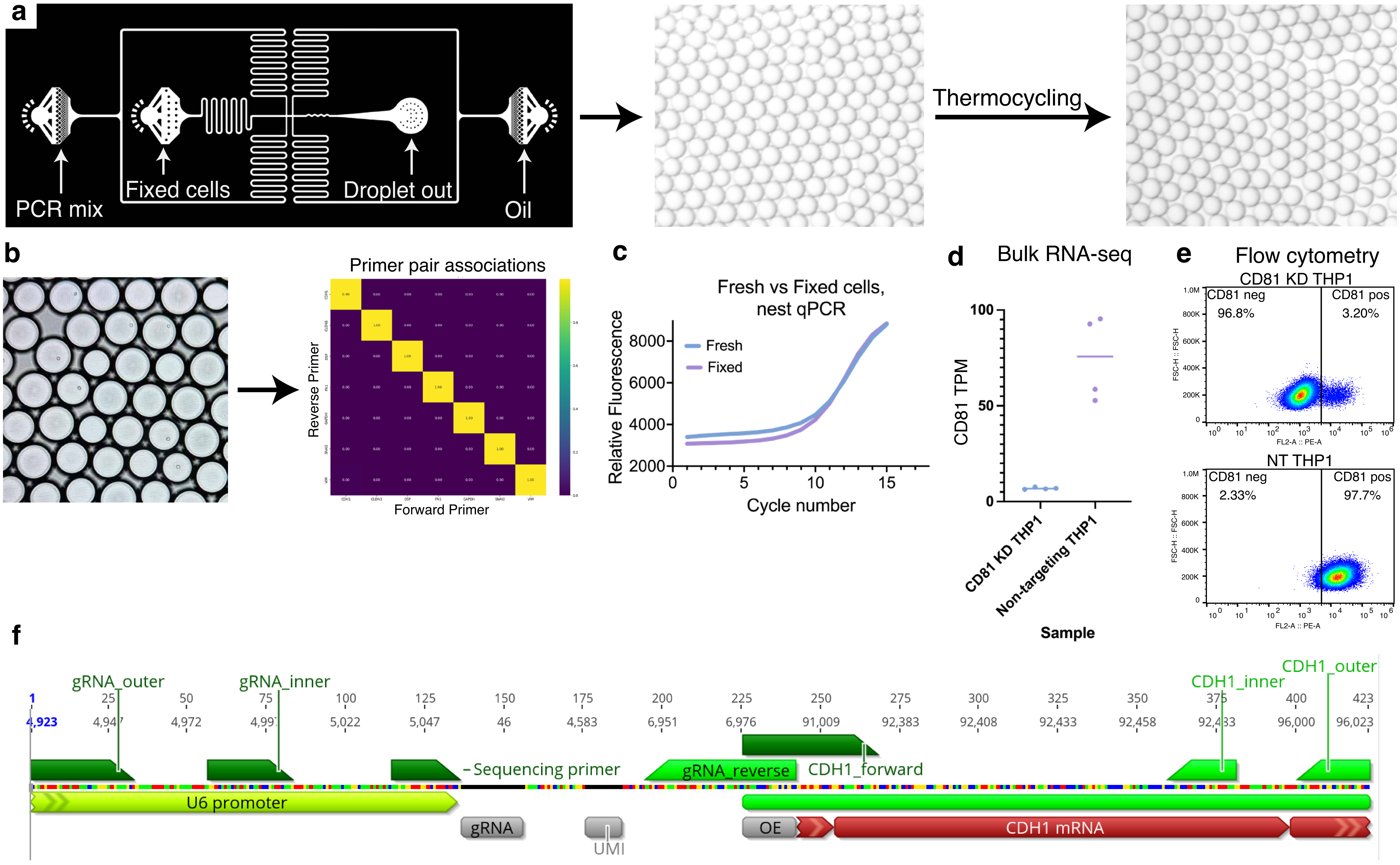

### Supplementary Figure 2

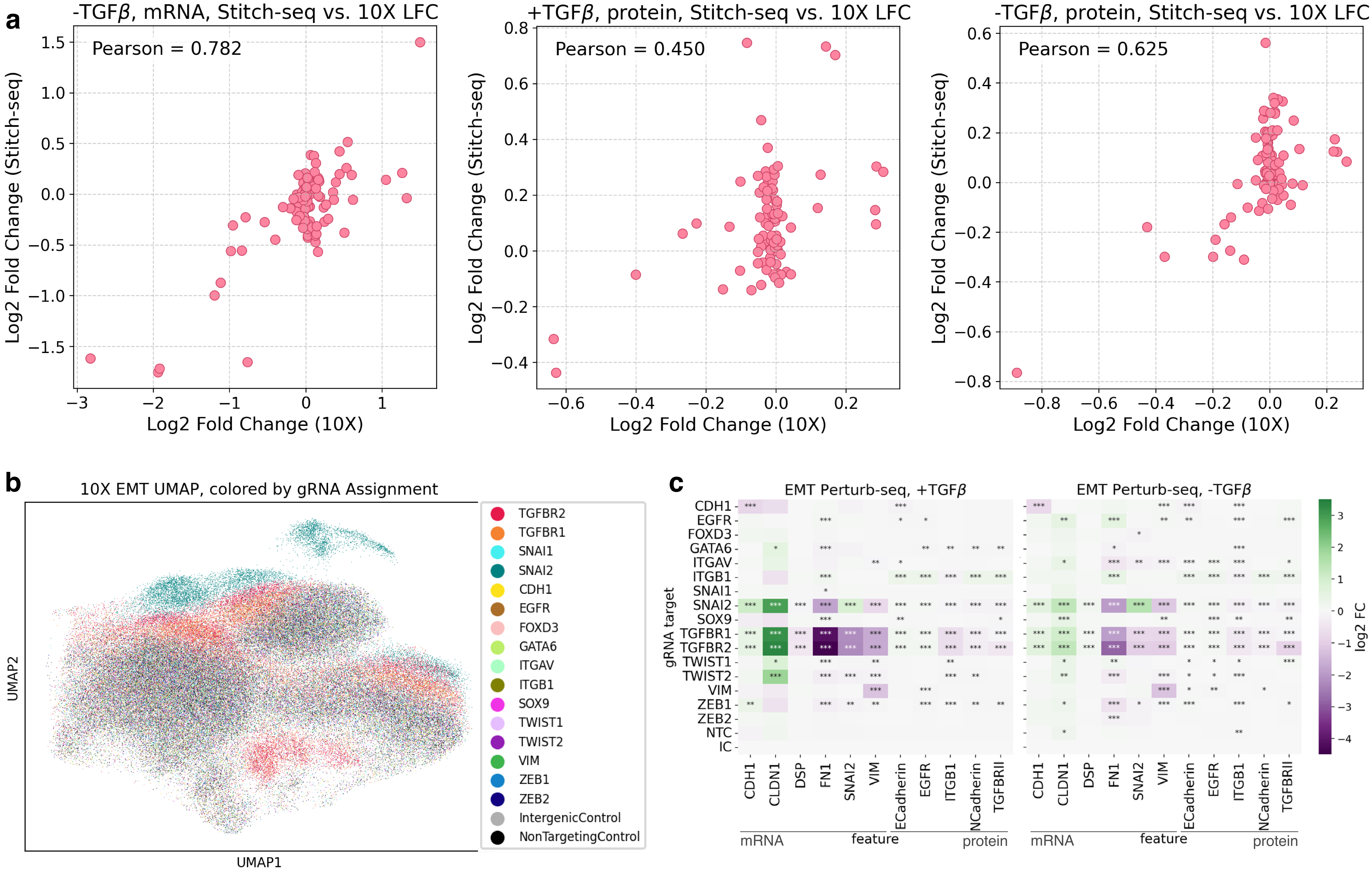

### Supplementary Figure 3

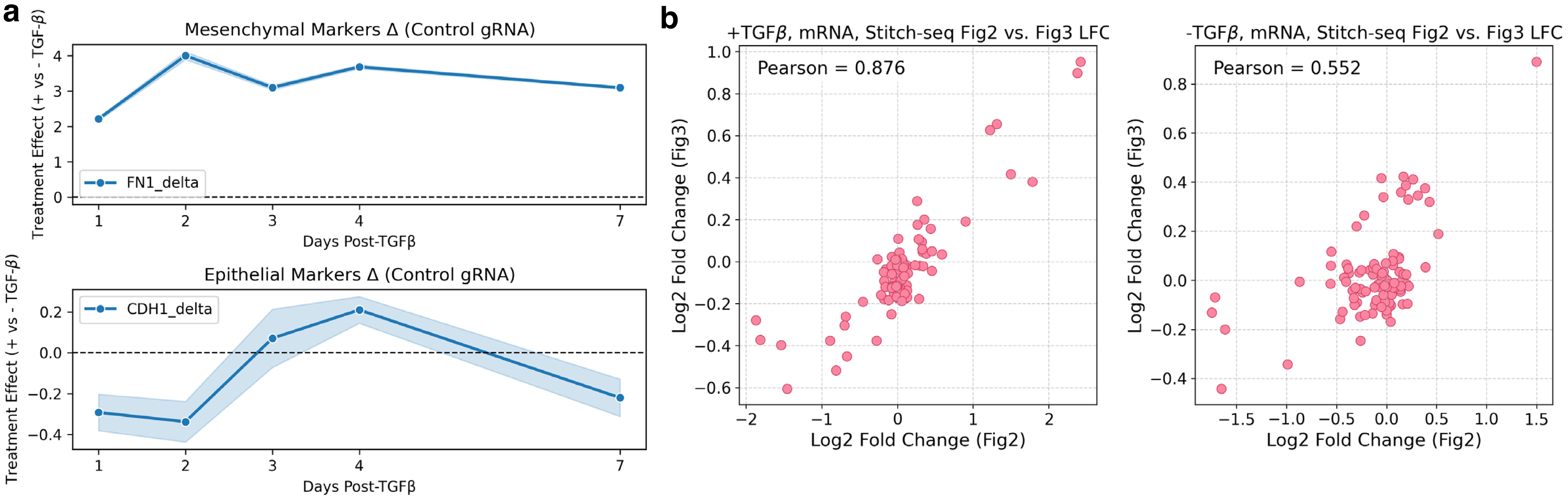
